# Borax-Based Gel Electrophoresis: A Novel Approach for RNA Integrity Analysis

**DOI:** 10.1101/2025.02.11.637722

**Authors:** Abdulhadi Albaser

## Abstract

This study explores the feasibility of using borax-based agarose gel electrophoresis for RNA integrity assessment. Multiple independent experiments demonstrated that borax, in addition to its buffering properties, exhibits denaturing-like behavior, effectively separating RNA molecules with comparable resolution to formaldehyde-based methods. This finding is surprising, given that theoretically, no evidence for borax denaturing activity exists, as it is typically considered a non-denaturing buffer. This method offers a compelling alternative to traditional formaldehyde-based methods for RNA analysis in various research and clinical settings, offering several advantages, including enhanced safety, simplified protocols, and reduced electrophoresis time. By eliminating the need for pre-treatment steps and utilizing borax as both a buffer and denaturant, this method simplifies RNA analysis. While the precise mechanism underlying this denaturing-like effect remains to be elucidated, further studies are needed to fully characterize these properties and explore its broader applicability.

## Introduction

Maintaining RNA integrity is crucial for accurate gene expression analysis, reliable RT-PCR results, and successful RNA sequencing [1, 2]. However, RNA is susceptible to degradation by RNases, ubiquitous enzymes found in various environments, including microorganisms, skin, and bodily fluids [3, 4, 5, 7]. These enzymes, present in viruses [1, 2], prokaryotes [3], eukaryotes [4, 5, 7], and other organisms, pose a significant threat to RNA integrity in research and clinical settings [8]. Traditional methods for assessing RNA quality often rely on hazardous chemicals such as formaldehyde and mercury hydroxide [13-16], posing significant safety risks and limiting accessibility. Furthermore, automated instruments, while effective, can be expensive and inaccessible to many researchers.

This study presents a novel approach for RNA quality assessment using borax-based agarose gel electrophoresis. Borax, a readily available and cost-effective buffer, offers several advantages over traditional methods, including enhanced safety by eliminating the need for hazardous chemicals like formaldehyde. The simplified protocol, which involves direct loading of RNA samples onto the gel without pre-treatment, reduces the time and effort required for RNA analysis. This method also offers the potential for reduced costs compared to some commercial reagents. This study demonstrates the feasibility of using borax as a buffer for RNA electrophoresis, enabling rapid and efficient assessment of RNA integrity.

## Methods

### Gel Preparation

- A 1% agarose gel was prepared using a 5 mM sodium tetraborate decahydrate (borax) solution as the running buffer.
- For comparison, a 1% agarose gel with standard TAE buffer was also prepared.
- Gel Red or ethidium bromide was added to both melted agarose solutions before casting.

### Sample Preparation

- RNA samples from *E. coli* were mixed with 1x colorless loading dye, 10X loading buffer containing glycerol (30%), EDTA (10 mM), SDS (0.5%), and water to 10 mL.

### Impact of Loading Buffer Components on RNA

- RNA samples were mixed with glycerol containing either SDS, EDTA, or glycerol only.
- Bromophenol blue can be optionally added.

### Electrophoresis

- Approximately 20 µL of RNA sample mixed with 2 µL of 10X loading buffer was loaded onto a 10 cm x 10 cm, 1% agarose gel.
- Electrophoresis was carried out at 120 V for 25-30 minutes for borax gels or 60 V for 60 minutes for TAE gels in a submerged gel electrophoresis system at room temperature.

### Concentration Effect of Borax on RNA Integrity

- To investigate the concentration-dependent effects of borax on RNA integrity, RNA samples were treated with a range of borax concentrations (50 μM - 10 mM).
- The treated samples were then loaded onto either borax or TAE gels and electrophoresed as described above.

## Results and Discussion

### Traditional RNA Electrophoresis Methods

Traditional RNA electrophoresis methods often rely on formaldehyde as a denaturant, necessitating pre-treatment steps such as heating RNA samples with formaldehyde and subsequent cooling on ice. These steps are time-consuming and increase the risk of RNA degradation. Furthermore, formaldehyde is a hazardous chemical, posing potential health and environmental risks. This highlights the need for safer and more efficient alternatives.

### Borax as an Alternative Buffer

Borax, a readily available and cost-effective compound, was chosen as an alternative buffer due to its known buffering capacity and ease of handling. This study explores the feasibility of using borax as an alternative buffer for RNA electrophoresis, aiming to develop a safer and more efficient method for RNA analysis. Borax effectively separated RNA molecules, as evidenced by the clear bands observed for 16S, 23S, and 5S rRNA in Figure 1. This observation suggests that borax can effectively resolve RNA species, demonstrating its potential as a denaturant for RNA electrophoresis without the need for traditional denaturants like formaldehyde. The sharp and distinct bands indicate high-quality RNA, comparable to those obtained with traditional denaturing gels. These results demonstrate that borax serves as a viable alternative to formaldehyde for RNA electrophoresis.

**Fig 1.**
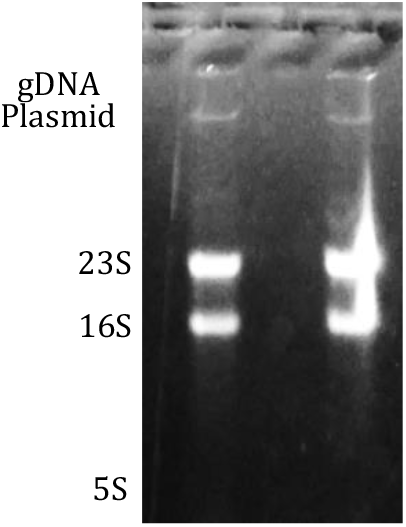
Separation of total *E. coli* RNA on borax agarose gel. The gel was prepared with 5 mM borax and run for 25 minutes at 120 volts. 16S and 23S rRNA bands, along with other RNA species (potential degradation products), were visualized after staining with ethidium bromide and exposure to UV transillumination.

### Alternative Approaches and Concentration-Dependent Effects

While formaldehyde is the standard denaturant for RNA electrophoresis, alternative approaches have been explored for RNA integrity assessment. For instance, Aranda and co-workers described a “bleach gel” approach using sodium hypochlorite for analyzing RNA quality [15]. This method has had a significant impact on the field, as evidenced by its high citation count. Sodium hypochlorite likely inactivates RNases, thereby preserving RNA integrity. Furthermore, hydrogen peroxide has also been explored as an alternative for RNA analysis in agarose gels [16]. Hydrogen peroxide may act as an oxidizing agent, potentially contributing to RNA stabilization.

To investigate the concentration-dependent effects of borax on RNA integrity, RNA samples were treated with a range of borax concentrations (50 μM - 10 mM). Higher concentrations of borax (2-10 mM) were found to completely degrade RNA (data not shown), likely due to its chaotropic properties at these elevated levels. However, lower concentrations (50-300 μM) did not significantly affect RNA integrity when run on TAE gels (Figure 3). The ratio of 2:1 is much clearer in TAE gel where samples were treated with different concentrations of borax, while the well separation of RNA bands is clearer in borax gel. This suggests that borax can effectively separate RNA molecules at lower concentrations without compromising RNA integrity.

### Unexpected Findings and Loading Buffer Components

Critically, and quite unexpectedly, clear separation of RNA bands was observed in the borax gel (Figure 2, Lane 1) when RNA samples were loaded with only glycerol as the loading buffer component, without any pre-heating or denaturation steps. This initial observation strongly suggested that borax itself might possess denaturing properties. Subsequent experiments confirmed this observation, as separation was achieved even in the absence of any additional denaturants or pre-treatment steps. Further experiments demonstrated that borax induces RNA separation in a manner similar to denaturing agents, as separation was observed in all tested components (Figure 2). These experiments were repeated several times with fresh buffers and fresh RNA extracts, yielding the same results. This is despite no previous studies indicating that borax may aid in denaturing RNA. This finding suggests that the conventional RNA electrophoresis protocols, often involving formaldehyde and pre-treatment steps such as heating and cooling, may not be necessary when using borax. Notably, with borax, any single loading buffer component, such as glycerol, can be used without the need for other components like EDTA, simplifying the sample preparation process.

**Fig 2.**
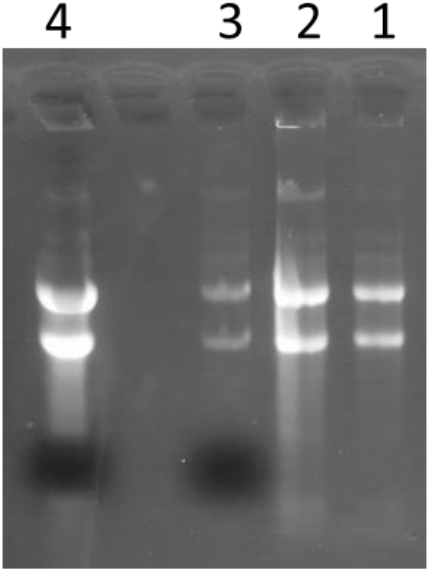
Effect of Loading Buffer Components on RNA Separation in Borax Agarose Gel. The gel was run for 25 minutes at 120 volts and stained with ethidium bromide. • Lane 1: RNA plus glycerol only. • Lane 2: RNA plus colorless loading buffer. • Lanes 3 & 4: RNA plus loading buffer containing bromophenol blue.

**Fig 3.**
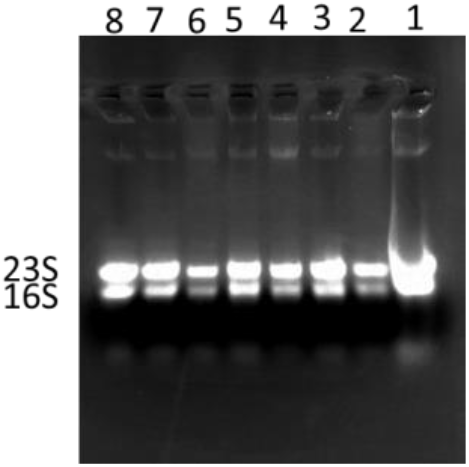
Effect of Borax Concentration on RNA Integrity. RNA samples were treated with varying concentrations of borax (50 μM - 300 μM) and subjected to electrophoresis (60 volts for 60 min) on TAE gels. • Lanes 1 and 8: Controls without borax. • Lanes 2-7: RNA samples treated with different borax concentrations.

### Mechanism of RNA Separation in Borax Gels

While the precise mechanism of RNA separation in borax gels requires further investigation, these findings demonstrate that borax contributes to RNA separation by maintaining a stable pH and ionic environment within the gel matrix, thereby minimizing RNA aggregation and promoting uniform migration, facilitating optimal RNA migration. The successful subcloning of plasmid DNA extracted from a borax gel further demonstrates the compatibility of this method with downstream applications.

### Future Research

While this study focuses on RNA integrity assessment, future research could explore the potential of borax-based gels for other applications, such as RNA extraction and purification. This study is primarily focused on *E. coli* RNA, and further research is needed to explore the applicability of this method to other RNA sources and to fully elucidate the mechanism of borax’s denaturing-like activity. Furthermore, investigating the long-term stability of RNA in borax gels would be valuable. Finally, a direct comparison of RNA integrity assessed by borax gels and traditional methods (e.g., Agilent Bioanalyzer) and investigating the compatibility of borax gels with different RNA isolation methods would strengthen the validation of this novel approach.

## Conclusion

This study demonstrates the feasibility of using borax as a buffer for RNA electrophoresis, offering a potential alternative to traditional formaldehyde-based methods. The borax-based method presents several advantages, including enhanced safety, a simplified protocol, and reduced electrophoresis time. While further research is needed to fully understand the underlying mechanisms, these preliminary findings suggest that borax-based gels can be a valuable tool for RNA integrity assessment in various research and clinical settings. By eliminating the need for hazardous chemicals and pre-treatment steps, the borax-based method enhances safety and efficiency, making it a more accessible technique for RNA analysis. Future studies will be essential to further validate this approach and explore its broader applications in RNA research and diagnostics.

## DATA AVAILABILITY STATEMENT

The raw data supporting the findings of this study are available within the article itself. No additional data are available.

## References

1. Taddeo, B., Maria Teresa Sciortino, Zhang, W., & Roizman, B. (2007). Interaction of herpes simplex virus RNase with VP16 and VP22 is required for the accumulation of the protein but not for accumulation of mRNA. Proceedings of the National Academy of Sciences of the United States of America, 104(29), 12163–12168.

2. Lim, D., Gregorio, G.G., Bingman, C., Martinez-Hackert, E., Hendrickson, W.A., & Goff, S.P. (2006). Crystal Structure of the Moloney Murine Leukemia Virus RNase H Domain. Journal of Virology, 80(17), 8379–8389.

3. Kazantsev, A.V., & Pace, N.R. (2006). Bacterial RNase P: a new view of an ancient enzyme. Nature Reviews Microbiology, 4(10), 729–740.

4. Jin, L., Kryukov, K., Suzuki, Y., Imanishi, T., Ikeo, K., & Gojobori, T. (2009). The evolutionary study of small RNA-directed gene silencing pathways by investigating RNase III enzymes. Gene, 435(1-2), 1–8.

5. Vieira, J., Fonseca, N.A., & Vieira, C.P. (2009). RNase-Based Gametophytic Self-Incompatibility Evolution: Questioning the Hypothesis of Multiple Independent Recruitments of the S-Pollen Gene. Journal of Molecular Evolution, 69(1), 32–41.

6. Brody, J.R., Calhoun, E.S., Gallmeier, E., Creavalle, T.D., & Kern, S.E. (2004). Ultra-fast high-resolution agarose electrophoresis of DNA and RNA using low-molarity conductive media. BioTechniques, 37(4), 598–602. 10.2144/04374ST04

7. Barber, G.N. (2009). The NFAR’s (Nuclear Factors Associated with dsRNA): Evolutionarily conserved members of the dsRNA binding protein family. RNA Biology, 6(1), 35–39.

8. Harder, J., & Schröder, J.M. (2002). RNase 7, a Novel Innate Immune Defense Antimicrobial Protein of Healthy Human Skin. Journal of Biological Chemistry, 277(48), 46779–46784.

9. Crooke, S.T., Lemonidis, K.M., Neilson, L., Griffey, R., Lesnik, E.A., & Monia, B.P. (1995). Kinetic characteristics of Escherichia coli RNase H1: cleavage of various antisense oligonucleotide-RNA duplexes. Biochemical Journal, 312(2), 599–608.

10. Bartholeyns, J., & Baudhuin, P. (1977). Purification of rat liver particulate neutral ribonuclease and comparison of properties with pancreas and serum ribonucleases. Biochemical Journal, 164(3), 675–683.

11. Dalaly, B.K., Eitenmiller, R.R., Friend, B.A., & Shahani, K.M. (1980). Human milk ribonuclease. Biochimica et Biophysica Acta (BBA) - Enzymology, 615(2), 381–391.

12. Kalnitsky, G., & Resnick, H. (1959). The Effect of an Altered Secondary Structure on Ribonuclease Activity. Journal of Biological Chemistry, 234(7), 1714–1717.

13. Kunitz, M. (1940). CRYSTALLINE RIBONUCLEASE. Journal of General Physiology, 24(1), 15–32.

14. Masek, T., Vopalensky, V., Suchomelova, P., & Pospisek, M. (2005). Denaturing RNA electrophoresis in TAE agarose gels. Analytical Biochemistry, 336(1), 46–50.

15. Aranda, P.S., LaJoie, D.M., & Jorcyk, C.L. (2012). Bleach gel: a simple agarose gel for analyzing RNA quality. Electrophoresis, 33(2), 366–369.

16. Pandey, R., & Saluja, D. (2017). Hydrogen peroxide agarose gels for electrophoretic analysis of RNA. Analytical Biochemistry, 534, 24–27.

17. Brody, J.R., & Kern, S.E. (2004). Sodium boric acid: a Tris-free, cooler conductive medium for DNA electrophoresis. BioTechniques, 36(2), 214–216.

